# The avian and human influenza A virus receptors sialic acid (SA)-α2,3 and SA-α2,6 are widely expressed in the bovine mammary gland

**DOI:** 10.1101/2024.05.03.592326

**Authors:** Charlotte Kristensen, Henrik E. Jensen, Ramona Trebbien, Richard J. Webby, Lars E. Larsen

## Abstract

An outbreak of H5N1 highly pathogenic influenza A virus (HPIAV) has been detected in dairy cows in the United States. Influenza A virus (IAV) is a negative-sense, single-stranded, RNA virus that has not previously been associated with widespread infection in cattle. As such, cattle are an extremely under-studied domestic IAV host species. IAV receptors on host cells are sialic acids (SAs) that are bound to galactose in either an α2,3 or α2,6 linkage. Human IAVs preferentially bind SA-α2,6 (human receptor), whereas avian IAVs have a preference for α2,3 (avian receptor). The avian receptor can further be divided into two receptors: IAVs isolated from chickens generally bind more tightly to SA-α2,3-Gal-β1,4 (chicken receptor), whereas IAVs isolated from duck to SA-α2,3-Gal-β1,3 (duck receptor). We found all receptors were expressed, to a different degree, in the mammary gland, respiratory tract, and cerebrum of beef and/or dairy cattle. The duck and human IAV receptors were widely expressed in the bovine mammary gland, whereas the chicken receptor dominated the respiratory tract. In general, only a low expression of IAV receptors was observed in the neurons of the cerebrum. These results provide a mechanistic rationale for the high levels of H5N1 virus reported in infected bovine milk and show cattle have the potential to act as a mixing vessel for novel IAV generation.

**Graphical abstract:** Created with Biorender.com

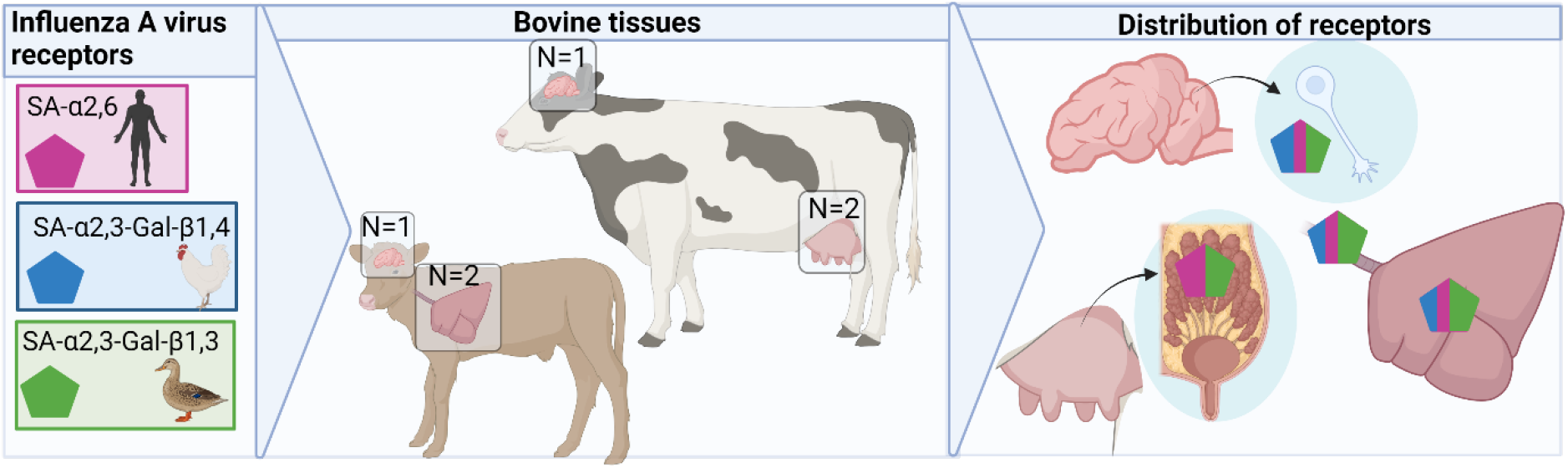

## 1. Introduction

The natural reservoir hosts of the influenza A virus (IAV) are waterfowls (Anseriformes) and shorebirds (Charadriiformes)^1^. IAV is a negative, single-stranded, RNA virus and, viral evolution has enabled some IAV to cross species barriers to establish in humans and a variety of mammals including pigs, horses, dogs, and seals^1,2,3^. Puzzlingly, cattle have until now been regarded almost resistant to infection with IAV, but susceptible to infection with influenza C and D viruses^4,5^. It was, therefore, with some surprise that highly pathogenic avian influenza virus (HPAIV) H5N1 (clade 2.3.4.4b) was detected in dairy cattle in Texas and rapidly spread to more than 40 herds in eight different states in the United States^6,7^. The clinical signs are dominated by sudden drop in milk production and mastitis, whereas only mild respiratory signs are observed and neurological symptoms that have often been described in other mammals infected by 2.3.4.4b viruses are absent^6,8^. A press release on April 25^th^ from the US Food and Drug Administration showed that one out of five retail milk samples tested positive for HPAI H5N1 by quantitative polymerase chain reaction (qPCR). These results combined with reports of detection of extremely high levels of virus in milk from infected cows^9^ in contrast to previous studies suggesting bovine milk inactivates the hemagglutinin (HA) of influenza viruses^5^. Previous studies have also shown a productive infection can be induced by the installation of a human influenza A isolate into the mammary glands of cows and goats^10,11^. Together, these findings indicate that the pathogenesis of HPAI in cattle differs from other mammals, thus, there is an urgent need for a better understanding of the pathogenesis of IAV in cattle and the anatomic features linking virus replication to mammary tissue.

HA binds to sialic acids (SA) terminally attached to glycans facilitating viral endocytosis and membrane fusion. One of the most well-described influenza virus species barriers is that the HA of human and swine adapted IAVs frequently prefer SAs linked to galactose (Gal) in an α2,6 linkage (SA-α2,6, human receptor), whereas avian IAVs prefer an α2,3 linkage (SA-α2,3, avian receptor)^12^. Furthermore, IAVs adapted to chickens generally prefer SA-α2,3-Gal with a β1,4 linkage to N-acetylgalactosamine (GalNac, SA-α2,3-Gal-β1,4-GalNac, referred to as the chicken receptor), whereas IAVs isolated from ducks favor SA-α2,3-Gal with a β1,3 linkage to N-acetylglucosamine (GlcNac, SA-α2,3-Gal-β1,3-GlcNac referred to as duck receptor)^13,14^. A study from 2011 investigated IAV receptor distribution on tracheal and lung tissues in cattle from Thailand and a more recent investigation assessed the receptor distribution on bovine primary cells of the nose, soft palate and trachea^15,16^. IAV receptor distribution in other bovine tissues are lacking.

Mass spectrometry can be used to examine the distribution of IAV receptors in tissues, however, *in situ* techniques are required to study the localization of the receptors in different cell types^17^. The Sambucus Nigra Lectin (SNA) binds to the human receptor, while Maackia Amurensis Lectin I (MAA-I) binds to the chicken receptor^18,19^. Maackia Amurensis Lectin II (MAA-II) shows a higher binding avidity for the duck receptor than the chicken receptor^20,21^. The main aim of this study was to investigate the *in situ* expression of IAV receptors in the bovine respiratory tract, cerebrum, and mammary glands by lectin histochemistry.

## 2. Materials and Methods

### 2.1. Tissue origination

Achieved bovine tracheal and lung tissues originating from two beef calves (2 months of age) were included. One of the calves showed acute, suppurative tracheitis. The following tissues were collected during routine necropsy from different clinical cases at the section of Veterinary Pathology, University of Copenhagen, Denmark. From a lactating dairy cow (4 years of age) two non-diseased mammary glands were included. The specimens from the cerebrum originated from one beef calf (5 months of age) and one dairy cow (2.5 years of age). The tissues were formalin-fixed, paraffin-embedded, and cut into 4-5 µm sections.

### 2.2. Lectin histochemistry

Detection of SA-α2,6 was performed using biotinylated SNA (B-1305-2, Vector Laboratories, California, USA) and detection of SA-α2,3 was performed using biotinylated MAA-1 (B-1315-2, Vector Laboratories) and biotinylated MAA-2 (B-1265-1, Vector Laboratories) as previously described^22^. The staining on the surface of the cells was evaluated as follows: -: no staining observed, 1: present in <50% of the cells, and 2: present in >50% of the cells.

## 3. Results

In the mammary gland, the human receptor (detected by the SNA lectin) and the duck receptor (detected by the MAA-II lectin) were widely distributed in the alveoli, but not in the ducts, whereas no positive staining of the chicken receptor (detected by the MAA-I lectin) was detected (Figure 1).

**Figure 1.**
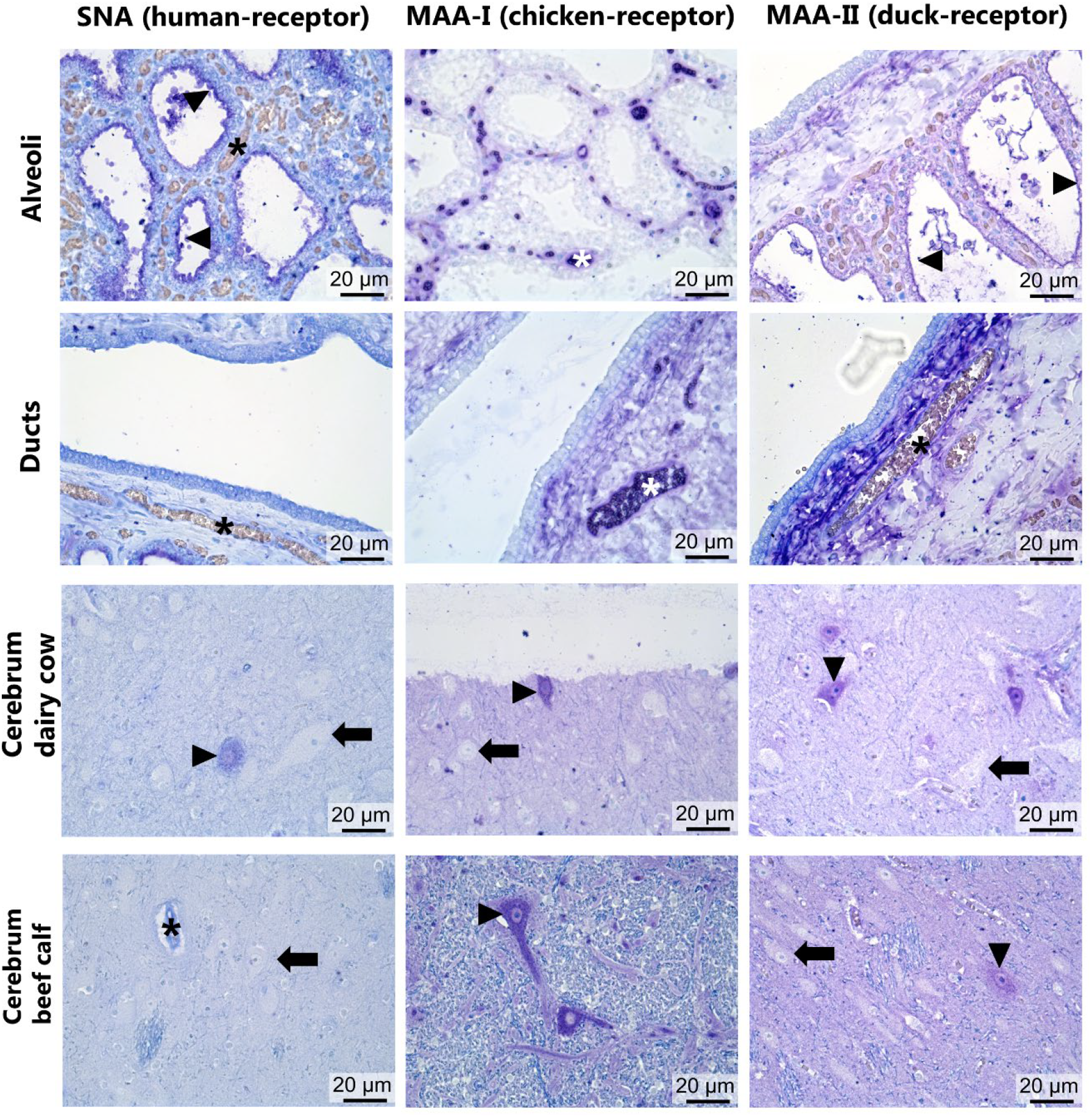
The distribution of human-, chicken- and duck influenza A virus receptors in the mammary gland tissue (alveoli and ducts) and in the cerebrum of cattle. The human-, chicken- and avian receptors was detected by Sambucus Nigra Lectin (SNA), Maackia Amurensis Lectin I (MAA-I), and Maackia Amurensis Lectin II (MAA-II) lectins, respectively, and positive reaction (dark blue to purple) was developed by adding Vector Blue. The arrowheads indicate positive staining of the surface epithelium in the udder tissue and cerebellar neurons, arrows indicate negative neurons, and asterisks indicate blood vessels.

In the respiratory tract, all receptors were expressed in the tracheal goblet cells but with a lower abundance of human receptor positive cells (Figure 2). Interestingly, only the chicken receptor was expressed on the surface of the ciliated epithelial cells (only observed in the calf with no tracheal inflammation). A sparse staining of the chicken receptor was expressed in the bronchi, and in the larger bronchioles, which also showed sparse staining for the duck receptor, and a scant staining of all three receptors was detected in the respiratory bronchioles. In the respiratory alveoli, all receptors were widely distributed but the human receptor appeared to have a specific affinity for the type II pneumocytes and/or leukocytes only.

**Figure 2.**
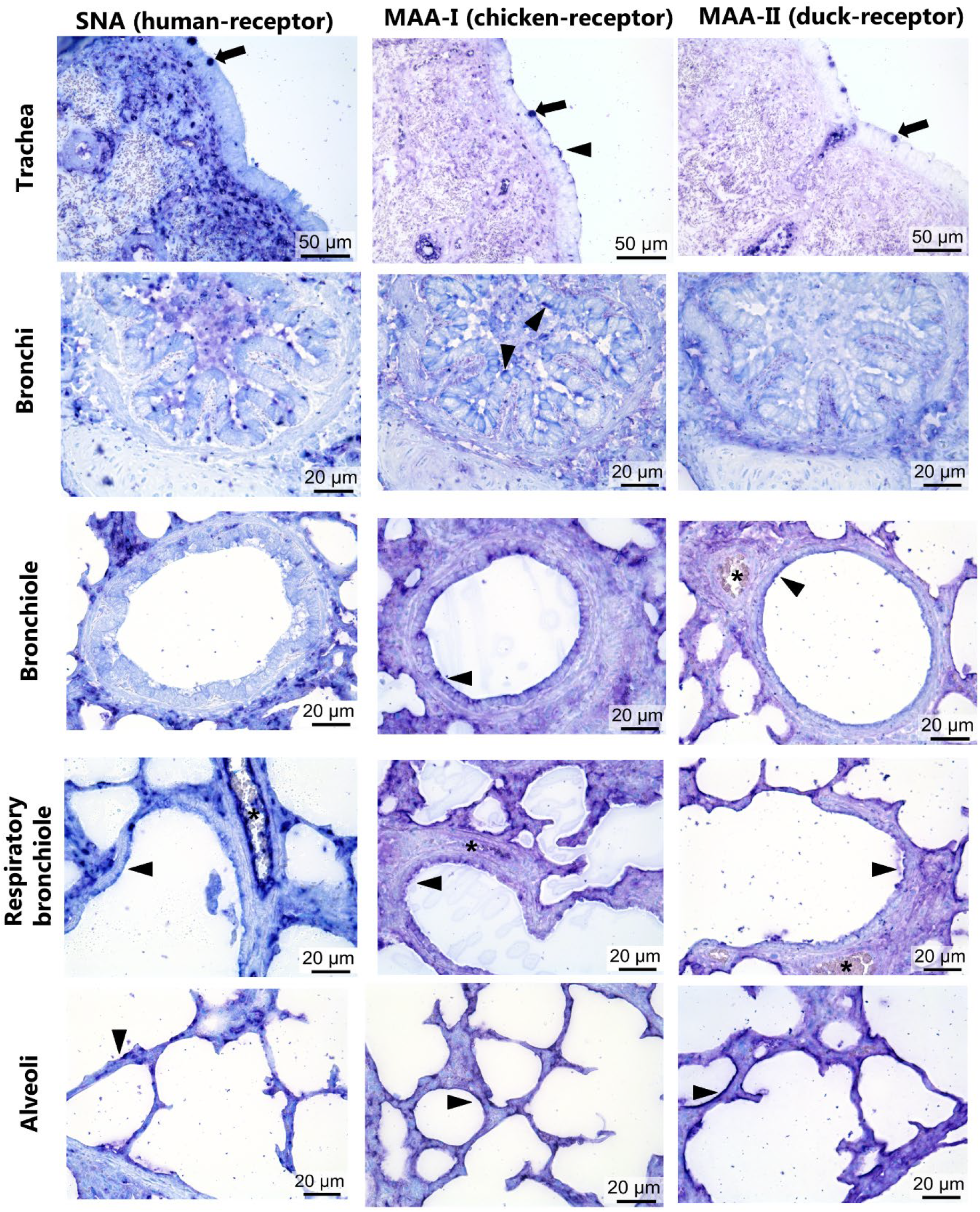
The distribution of human-, chicken- and avian influenza A virus receptors in the bovine respiratory tract. The human-, chicken- and avian receptors were detected by Sambucus Nigra Lectin (SNA), Maackia Amurensis Lectin I (MAA-I), and Maackia Amurensis Lectin II (MAA-II) lectins, respectively, and positive reaction (dark blue to purple) was developed by adding Vector Blue. The arrowheads indicate positive staining of the surface epithelium and asterisks indicate blood vessels.

A few, low-intensity cerebellar neurons stained positive for all receptors in the dairy cow with the lowest number positive for the human-receptor. In the beef calf only the duck- and chicken receptors were detected (Figure 1).

Lastly, positive staining of the human and chicken receptors was detected in the endothelial cells and the chicken receptor was also detected in the bovine erythrocytes. The results from the lectin histochemistry are found in Table 1 and summarized in Figure 3.

**Table 1.** The comparison of the distribution of influenza A virus (IAV) based on the semi-quantification of the lectin histochmistry.

| **Tissue** | **Cell type** | **SNA**<br>**(human-receptor)** | **MAA-I**<br>**(chicken-receptor)** | **MAA-II**<br>**(duck-receptor)** |
| --- | --- | --- | --- | --- |
| **Trachea** | Ciliated epithelium | - | 1<sup>1</sup> | - |
|  | Goblet cells | 1 | 2<sup>1</sup> | 2 |
|  | Glands | -<sup>2</sup> | 2 | 2 |
| **Bronchi** | Epithelium | -<sup>2</sup> | 1 | - |
| **Bronchiole** | Epithelium | - | 1 | 1 |
| **Respiratory bronchiole** | Epithelium | 1 | 1 | 1 |
| **Lung parenchyma** | Alveoli | 2<sup>3</sup> | 2 | 2 |
| **Cerebrum (dairy cow)** | Neurons | 1 | 1 | 1 |
| **Cerebrum (beef calf)** | Neurons | - | 1 | 1 |
| **Mammary gland** | Ducts | - | - | - |
|  | Alveoli | 2 | - | 2 |

-: no staining, 1: staining observed in <50% of the cells, 2: staining observed in >50% of the cells.
<sup>1</sup>The values was observed for the one beef calf without any lesions present, whereas the beef calf with acute, suppurative tracheitis showed markedly reduced staining than mentioned in the table.
<sup>2</sup>Positive staining observed in the lumen but not on the surface of the cells.
<sup>3</sup>Presumably type II pneumocytes or leukocytes.

**Figure 3.**
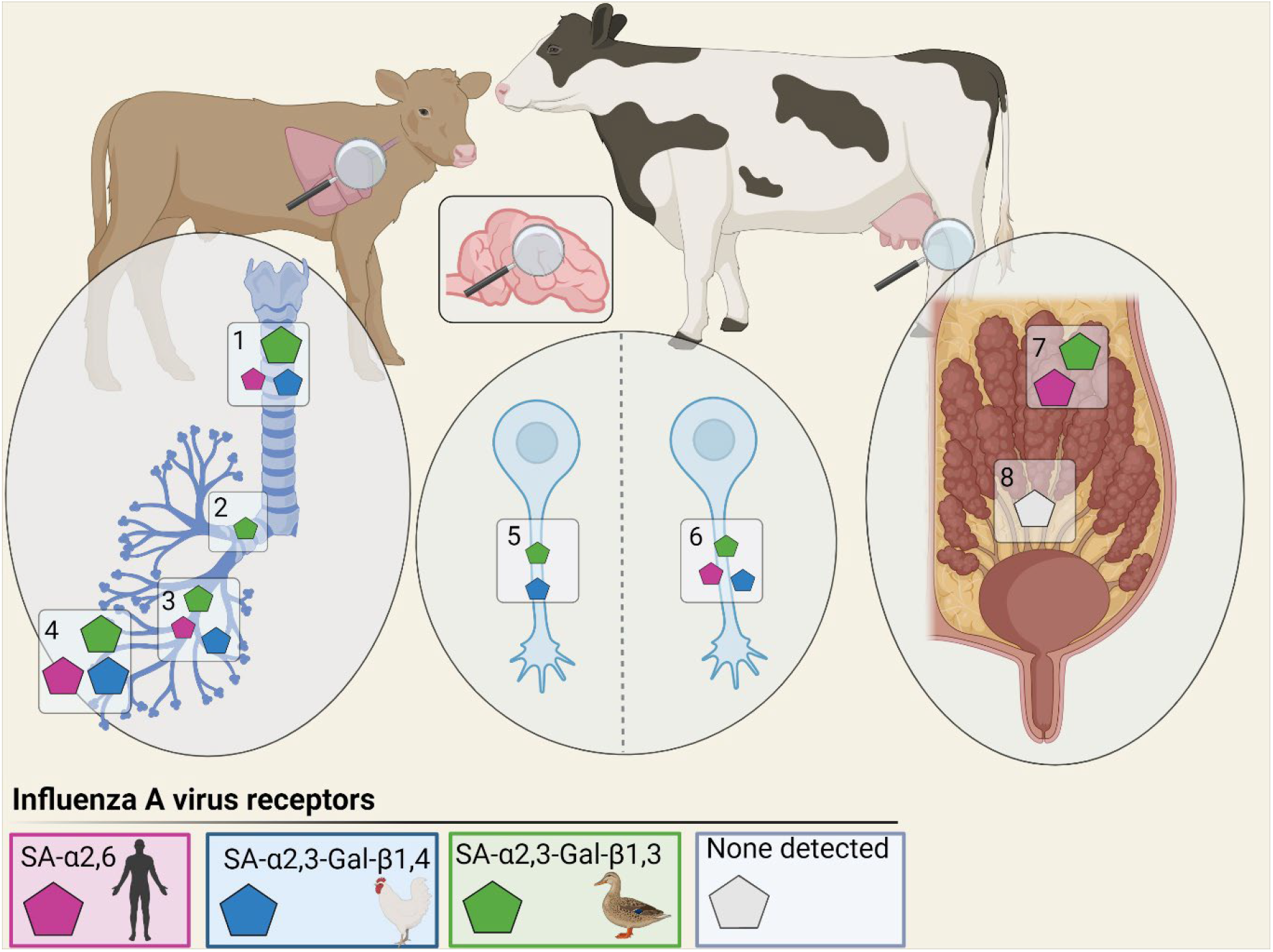
Comparison of the distribution of influenza A virus (IAV) receptors in the lungs, cerebrum, and mammary gland tissues of cattle. The distribution and staining intensity of the IAV receptors was detected by lectin histochemistry and semi-quantified. The human receptor (pink) was detected by Sambucus Nigra Lectin (SNA), the chicken receptor (blue) was detected by Maackia Amurensis Lectin I (MAA-I), and the duck receptor (green) was detected by Maackia Amurensis Lectin II (MAA-II). The tissues investigated were the trachea (1), bronchi (2), bronchioles (3) alveoli (4) of the bovine respiratory tract, and the cerebellar cortex of a beef calf (5) and a dairy cow (6) and lastly the alveoli (7) and ducts (8) of the mammary gland. Created with Biorender.com.

## 4. Discussion

Here we evaluate the expression of IAV receptors *in situ* in the mammary gland, respiratory tract and cerebrum of cattle, which typically has been considered less susceptible to IAV infection^5^. Strikingly, was the finding that both the human- (SA-α2,6) and the duck receptors (SA-α2,3-Gal-β1,3) were highly expressed in the mammary glands, whereas no expression of the chicken receptor (SA-α2,3-Gal-β1,4) was detected. A previous study showed that co-expression of both the human- and avian receptors can enhance the receptor binding of H5N1 isolated from ducks (clade 2.1.1) *in vitro*^23^. Combined these findings support the hypothesis that the high viral load seen in milk from cows infected by HPAI H5N1 virus belonging to clade 2.3.4.4b are due to local viral replication, because these viruses have high affinity for this receptor^24^. Additionally, the avian receptor has been found to be highly expressed in the human cornea and conjunctiva^25^ which may explain the report that conjunctivitis was the dominating clinical sign of a person presumably infected by dairy cows in Texas^7^.

The transmission route(s) and the pathogenesis of H5N1 in cows remain unclear, and it’s not known if the virus enters the mammary gland by an ascending infection or systemically by the blood supply. Interestingly, neither the human-, chicken-, nor the duck receptors were expressed in the ducts of the mammary gland, making an ascending mammary gland infection more challenging. It is not clear to which degree the HPAIV infected cows develop viremia, however, even a very low degree of viremia may be adequate for the virus to enter the mammary gland and establish infection because the blood flow in the lactation period is ∼400 liter per hour^26^. Suggestions by the USDA that only some udder quarters may be involved in infection does, however, argue against a viremic source^27^.

The investigation of the IAV receptor distribution in the respiratory tract also revealed some novel findings. In the upper respiratory tract and upper part of the lower respiratory tract (trachea, bronchi, and bronchioles), the chicken receptor (SA-α2,3-Gal-β1,4) was expressed on the surface of the respiratory epithelium, whereas a lack of - or very limited expression - of the human and duck receptors was detected. This pattern is the opposite to what we found in the mammary gland. The lack of expression of the human receptor in the upper respiratory tract of cattle contrasts with findings in humans^25,28^ and swine^22,25^ and supports the perception that bovines are highly resistant to infection with influenza A viruses of human and swine origin when exposed by the respiratory route^1,29^. In the lung alveolar cells, however, all three receptors were abundantly expressed, similar to what has been found in pigs and humans^22,25,28^.

Isolated primary respiratory cells from the nasal turbinates, soft palate, and tracheal tissues of cattle had an IAV receptor distribution that corresponds to the findings of our study^16^. Another previous study investigated the presence of the human receptor (SNA, SA-α2,6-Gal) and avian receptors (MAA, SA-α2,3-Gal) by lectin histochemistry in the bovine trachea and lung tissues, and found only the human receptor expressed in the lung tissues^15^. The discrepancy between results from this study^15^ and our results could be due to the lack of citrate pre-treatment of the tissues in the previous, which has been shown to markedly increase the staining of the lectins in formalin-fixed tissues^28^.

The HPAIV H5N1 virus (clade 2.3.4.4b) has a global distribution in wild birds and has infected more than 35 different species of mammals^24^. One of the hallmarks of most of these infections has been the dominance of neurological symptoms and post mortem have detected viruses in the brain of dead animals^30^. We therefore investigated the receptor expression in the cerebrum of cattle and found no or very scarce expression of any of the receptors. The sparse expression of the IAV receptor in the cerebrum of cattle may explain why there is a lack of neurological signs in HPAIV-infected cows, however, this remains speculative because the pathogenesis of the neurotropic HPAIV virus in other species remains unclear.

In conclusion, here we provide new insight into IAV host receptors in cattle. The expression of the duck receptor in the mammary gland of cows fits well with the observed widespread infections among cattle in the United States. Nevertheless, the presence of the IAV receptors does not per se provide evidence for cattle being susceptible to all avian influenza viruses. The co-expression of both human and avian receptors in the mammary glands indicate susceptibility for viruses of both swine/human and avian origin. This is worrying from a zoonotic perspective, because bovines may act as a mixing vessel for new IAVs with increased zoonotic potential.

Additional research is very much needed to better understand the pathogenesis and epidemiology of IAV infections of cattle and other ruminants to elucidate if these species can act as a mixing vessel for new IAVs.

## Acknowledgements

This work was supported by the Novo Nordic foundation (FluZooMark: NNF19OC0056326). The authors thank Elisabeth Wairimu Petersen and Betina Gjedsted Andersen for practical laboratory help. We declare that we have no conflict of interest.

